# Dysregulation of energy metabolism and calcium homeostasis in iPSC-derived neurons carrying Presenilin-1 M146L gene mutation

**DOI:** 10.1101/2025.09.08.674945

**Authors:** Carlos Wilson, Pablo Galeano, María Mónica Remedi, Gisela V Novack, Lorenzo Campanelli, Laura Gastaldi, Esteban Miglietta, Andrés H. Rossi, Natividad Olivar, Luis I Brusco, Eduardo M Castaño, Alfredo Cáceres, Laura Morelli

**Affiliations:** Centro de Investigación en Medicina Traslacional “Severo R Amuchástegui” (CIMETSA), Instituto Universitario Ciencias Biomédicas Córdoba (IUCBC), Córdoba, Argentina; Laboratory of Brain Aging and Neurodegeneration, Fundación Instituto Leloir, IIBBA-CONICET, Av. Patricias Argentinas 435, C1405BWE, CABA, Buenos Aires, Argentina; Microscopy and BioImaging Facility, Fundación Instituto Leloir-IIBBA-CONICET, Buenos Aires, Argentina, Av. Patricias Argentinas 435, C1405BWE, CABA, Buenos Aires, Argentina”; Center of Neuropsychiatry and Neurology of Behavior, School of Medicine, University of Buenos Aires, Buenos Aires, Argentina

**Author notes:** These authors contributed equally to this work and share senior authorship (Alfredo Cáceres, Laura Morelli).

**Keywords:** Familial Alzheimer’s disease, iPSC-derived neural stem cells, mitochondrial respiration, mitofusin, calcium dynamics, reactive oxygen species (ROS)

## Abstract

Impaired cellular activities, particularly in highly active cells such as neurons, are primarily supported by metabolic abnormalities and failures in Ca²⁺ homeostasis. Here, we provide an integrative analysis of human iPSC-derived neurons (iNs) carrying the Presenilin-1 M146L gene mutation (PS1^M146L^) and control cells (PS1^control^). PS1^M146L^ iNs exhibited abnormal Ca²⁺ dynamics, a significant increase in key parameters of mitochondrial respiration, and higher intracellular ROS levels. KCl-evoked depolarisation was significantly lower in PS1^M146L^, suggesting a failure in maintaining the electrochemical gradient across the plasma membrane. Following thapsigargin stimulation, mitochondrial Ca²⁺ levels ([Ca²⁺]m) were significantly reduced in PS1^M146L,^ while [Ca²⁺]m did not differ significantly between genotypes after treatment with bradykinin, suggesting that impairments in the [Ca²⁺]m homeostasis are particularly evident under stress conditions and do not impact the 1,4,5-triphosphate (IP3) pathway. Since iNs of both genotypes were sensitive to the MCU-1 inhibitor, the deficits observed in PS1^M146L^ could be the consequence of impairments in the ER-mitochondria contacts. Our results illustrate the utility of iNs carrying PS1 mutations in understanding how human neurons alter relevant pathways before neurodegeneration.

## Introduction

More than 180 Presenilin 1 (PS1) mutations, that are associated with familial Alzheimer’s disease (FAD), have been identified at 121 distinct sites (www.alzforum.org/mutations), yet the precise role of PS1 in Alzheimer’s pathology remains debated. PS1 is a transmembrane protein mainly localised in the secretory pathway (Annaert et al., 1999; Culvenor et al., 1997; Walter et al., 1996), with smaller amounts also detected at the cell surface (Ray et al., 1999). According to the amyloid hypothesis, PS1-linked mutations increase the production and relative abundance of amyloidogenic Aβ42 species, promoting aggregation and neuronal loss (Selkoe and Hardy, 2016). However, extensive in vitro studies examining Aβ42 and Aβ40 generation in the presence of multiple pathogenic PS1 mutations indicate that altered proteolytic activity alone is unlikely to fully account for FAD (Sun et al., 2017). Beyond its role in γ-secretase activity, PS1 has been involved in several non-catalytic functions, such as lysosomal acidification, autophagy, protein trafficking and Ca^2+^ regulation (Otto et al., 2016). Several studies demonstrate that PS1 modulates ER Ca^2+^ homeostasis. Studies performed in human fibroblasts and in primary neurons from transgenic mice demonstrated that PS1 interacts with the inositol 1,4,5-triphosphate receptor (IP3R) (Cheung et al., 2008) and with the sarco/endoplasmic reticulum Ca²⁺-ATPase (SERCA) (Green et al., 2008), indicating a direct role for PS1 in the regulation of intracellular Ca²⁺ handling. In addition, PS1 has been proposed to function as a ER-Ca^2+^ leak channel (Tu et al., 2006), and several FAD mutations impair this leak, leading to ER Ca^2+^ overload (Nelson et al., 2007). Nonetheless, the classification of PS1 as a Ca^2+^ channel remains controversial, as other groups have reported conflicting results (Cheung et al., 2010).

PS1 is highly enriched at ER-mitochondria associated membranes, a key subcellular site for calcium regulation (Hayashi et al., 2009). According to the “Ca^2+^ overload” hypothesis of FAD (LaFerla, 2002), PS1 mutations promote excessive ER Ca^2+^ release, which activates the mitochondrial calcium uniporter (MCU-1) and drives increased mitochondrial Ca^2+^ uptake. This, in turn, enhances mitochondrial respiration (Glancy and Balaban, 2012) and elevates reactive oxygen species (ROS) production. Such a mechanism is consistent with the view that mitochondrial dysfunction represents a central event in Alzheimer’s disease pathology (Swerdlow et al., 2010). Despite this, little is known about energy metabolism in living human neurons. For decades, Alzheimer’s disease models relied on immortalised cell lines, primary neuronal cultures from transgenic rodents, or fibroblasts from FAD patients, which limited insights into disease mechanisms in human neurons. The emergence of neurons derived (iNs) from human induced pluripotent stem cells (iPSCs) now enables the study of AD-related processes in a human and genetically defined context. Because clonal expansion reduces variability between cells, iNs are particularly valuable for dissecting the contribution of specific mutations to disease phenotypes (Gudenschwager et al., 2021). In this regard, recent studies reported that iNs carrying PS1 mutations (S290C or A246E) did not show alterations in Aβ42/Aβ40 ratios but displayed increased Ca^2+^ responses to glutamate or AMPA stimulation compared with their isogenic controls (Targa Dias Anastacio et al., 2024). These findings suggest that PS1 mutations perturb Ca^2+^ homeostasis, rendering neurons more sensitive to excitatory stimulation.

In this study, we examined iNs derived from a member of a previously described Argentine pedigree (AR1) carrying the PS1 M146L mutation (PS1^M146L^) (Morelli et al., 1998), along with iNs from a healthy individual (PS1^control^). We specifically analysed Aβ production, mitochondrial morphology and respiration, in situ ROS generation, and cytosolic and mitochondrial Ca^2+^ dynamics, both under basal conditions and after treatment with i11, a selective inhibitor of the mitochondrial Ca^2+^ uniporter (MCU-1). This iPSC-based model of FAD provides a valuable platform to investigate how neurons from mutation carriers alter key molecular pathways before the onset of overt neurodegeneration.

## Results

### Generation of a human cellular model to address the pathophysiological mechanisms of the PS1^M146L^ mutation

Dermal fibroblasts were collected at the Hospital de Clínicas of the School of Medicine of the University of Buenos Aires (Argentina) from 2 donnors: #1) a carrier (V-36) aged 35 years, belonging to the Argentine pedigree (ARl) (Fig. S1A) bearing the PS1^M146L^ gene mutation in which one deceased member showed neuropathological confirmation (Fig. S1B) and #2) a cognitively normal subject (not a mutant carrier and not related to subject #1), aged 39 years and referred to as _PS1_^control^.

To establish a cellular model that can inform on the pathophysiological mechanisms of PS1^M146L^, we reprogrammed a subset of fibroblasts into iPSCs (see Materials and Methods section). In this report, we assessed two different clones of each genotype, F2A112 and F2A121 for PS1^control^, and F22A111 and F22A23 for PS1^M146L^. Correct mutation propagation of A>T transversion in exon 5 was verified by Barcode-Tagged Sequencing™ (Celemics, Korea) in clones F22A23 and F22A111 (Table S1).

To assess amyloidogenic AβPP processing, the Aβ levels were measured in conditioned cell media. Compared with PS1^control^, γ-secretase activity in PS1^M146L^ iNs did not significantly alter the production of Aβ38 (1.1 µg/mL (IQR: 0.7-1.7) vs. 2.1 µg/mL (IQR: 0.7-4.1); U=12, p=n.s., Mann–Whitney test) or Aβ40 (11.4 ± 2.7 µg/mL vs. 13.5 ± 5.9 µg/mL; t=0.37, p=n.s.). Although not statistically significant, Aβ42 levels were higher in PS1^M146L^ iNs (1.4 µg/mL (IQR: 0.9-3.0) vs. 0.8 µg/mL (IQR: 0.6-1.9); U=9, p=n.s., Mann-Whitney test). Consistently, the Aβ42/40 ratio showed a non-significant trend towards an increase (1.7 (IQR: 1.5-2.4) vs. 0.6 (IQR: 0.4-2.0); U=6, p=0.09, Mann-Whitney test). In summary, iNs carrying the PS1^M146L^ mutation exhibited at *days in vitro* (DIV) 14 non-significant higher Aβ42 levels and a trend toward an increase in the Aβ42/40 ratio compared with PS1^control^. To establish a framework of analysis, we characterised the progression of neural differentiation by immunofluorescence and confocal imaging, using lineage-specific markers in control cells (Fig. 1). In particular, we assessed the proportions of neural progenitor cells (NPCs) and neurons within the first 21 DIV differentiation. Immunofluorescence staining was performed using antibodies against SOX2 (neuronal progenitors), Nestin (NSCs), βIII-Tubulin (immature neurons), and MAP2 (neurons) at 7, 14, and 21 DIV (Fig. 1A). As expected, the outcome of differentiating iNs was heterogeneous, showing a mixture of NPCs and post-mitotic neurons in culture (Fig. 1B). Nevertheless, βIII-tubulin and MAP2-positive populations were significantly enriched over time, shifting the identity of the culture from NPCs to neurons after 7 DIV.

**Figure 1.**
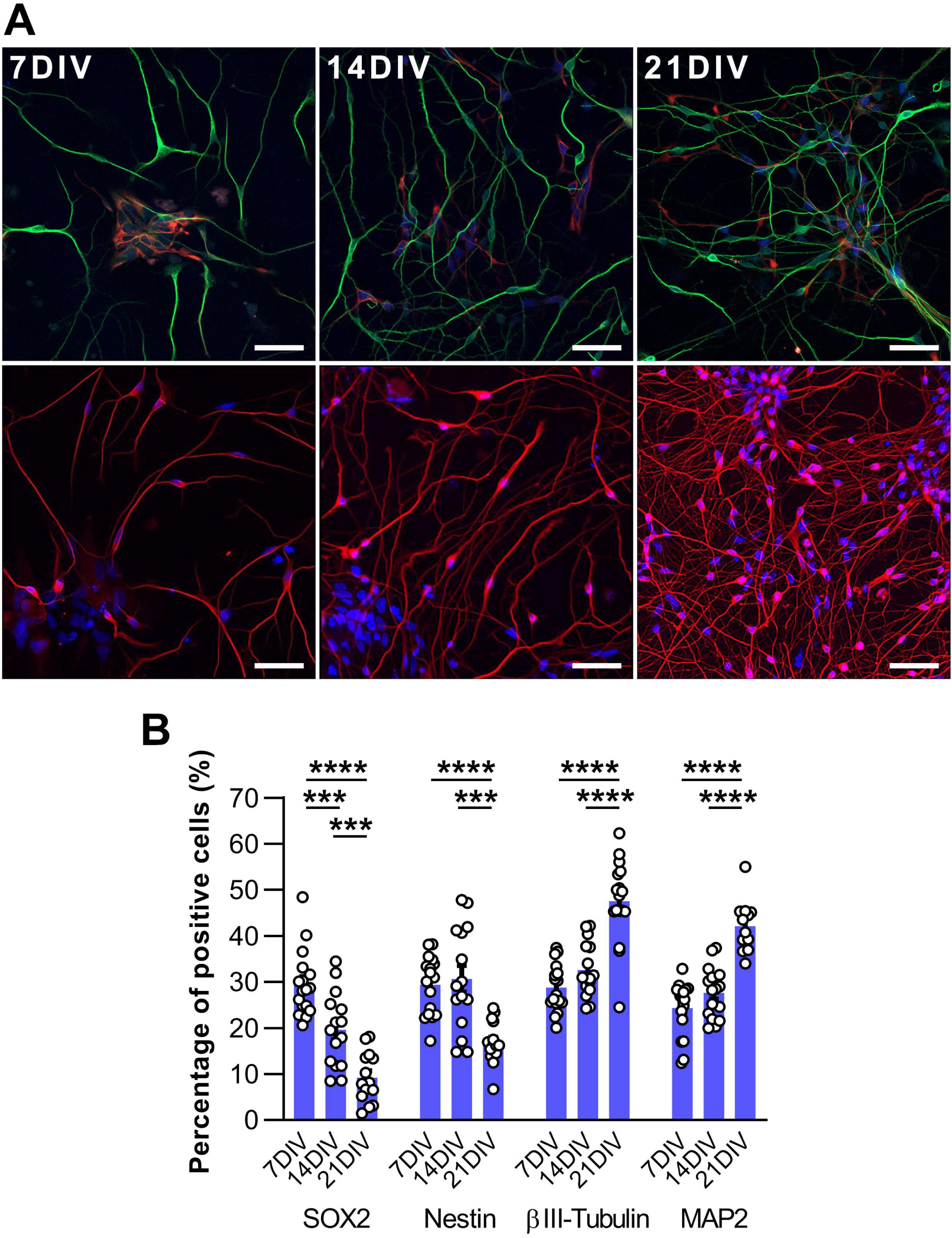
Morphological characterization of PS1^control^ iNs in culture. **(A)** Upper panel. Representative confocal images showing molecular markers MAP2 (green), nestin (red) and SOX2 (blue), identifying neurons, neural stem cells (NSCs), and neuronal progenitors, respectively, at 7, 14 and 21 DIV. Scale bars, 50 µm (20x); Lower panel. Representative images showing βIII-tubulin (red) and DAPI (blue) at 7, 14 and 21 DIV. Scale bars, 50 µm (20x). **(B)** Quantification of neuronal (βIII-tubulin, MAP2) and NSC populations (nestin, SOX2) using High Content Analysis (HCA) microscopy. Bars and error bars show the mean ± SEM with individual values superimposed. ***p<0.001; ****p<0.0001, Tukey’s (SOX2, βIII-tubulin, MAP2) or Games–Howell’s (nestin) multiple comparisons tests after significant one-way ANOVA or Welch’s ANOVA, respectively (SOX2: F=32.08, p<0.0001; nestin: W=25.09, p<0.0001; βIII-tubulin: F=35.54, p<0.0001; MAP2: F=36.72, p<0.0001).

To further confirm the progressive acquisition of neuronal properties, we evaluated the excitability of 7 to 21 DIV control iNs by live-cell Ca^2+^ imaging using the fluorescent reporter Fluo4-AM (Fig. 2). After 30 s of baseline recording, iNs were stimulated with 120 mM KCl to induce extracellular Ca^2+^ uptake as a readout of depolarisation. We were able to visualise a consistent KCl-evoked response in 14 and 21 DIV iNs (Fig. 2A, 2C, 2E). Whilst left plots show the mean trend, right plots show the cell-by-cell response, highlighting a heterogeneous behaviour. Particularly, at 7 DIV, most of the cells remained unresponsive, and just a few were able to uptake Ca^2+^ after KCl stimulation. By contrast, 14 and 21 DIV iNs were highly responsive, suggesting that iNs become progressively excitable over time in culture. Similarly, we evaluated the Ca^2+^ uptake in response to 200 μM glycine, a co-agonist of NMDAR (Johnson and Ascher, 1987; Kleckner and Dingledine, 1988; Guo et al., 2017). Accordingly, iNs showed a consistent uptake starting at 14 DIV, suggesting the onset of a glutamatergic phenotype, characteristic of brain cortical neurons (Fig. 2B, 2D, 2F). Together, Fig. 1 and 2 are instrumental to understanding the molecular and physiological properties of control iNs used in this work.

**Figure 2.**
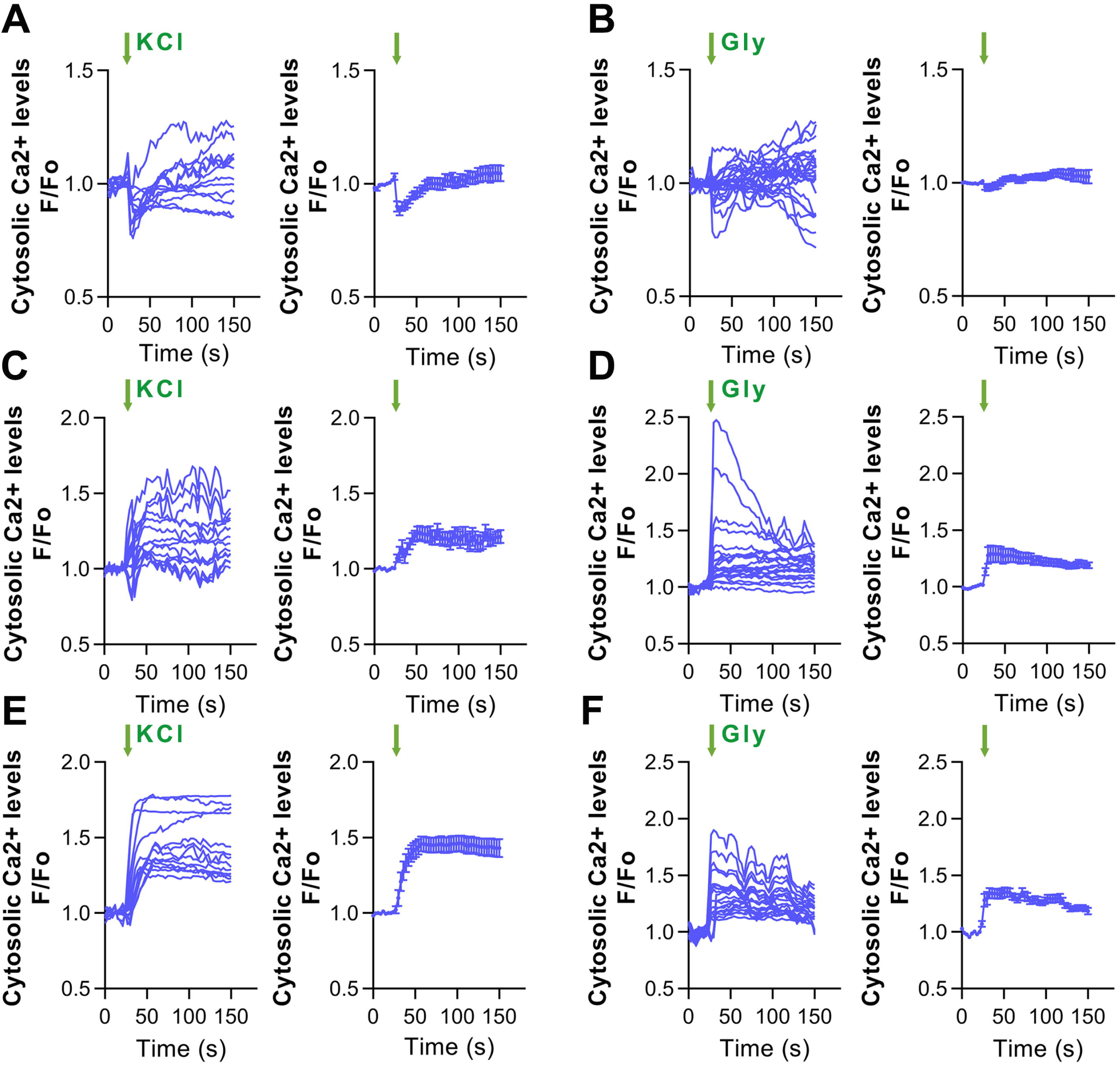
Analysis of excitability and glutamatergic properties of PS1^control^ iNs by Fluo4 Ca^2+^ imaging. (A, C, E). KCl-evoked Ca2+ entry in 7, 14, and 21 DIV PS1^control^ iNs (respectively), using Fluo4 and live-cell Ca^2+^ imaging. Graphs represent the time-lapse response of KCl-evoked Ca^2+^ entry (left panels: individual values; right panels: mean data ± SEM). **(B, D, F)** Glycine-evoked Ca^2+^ entry in 7, 14, and 21 DIV PS1^control^ iNs (respectively), using Fluo4 and live-cell Ca^2+^ imaging. Graphs represent time-lapses of glycine (Gly)-evoked Ca^2+^ entry (left panels: individual values; right panels: mean data ± SEM). N=3 biological replicates.

Moreover, we assessed the ability of neural cells to undergo polarisation— used as an indicator of neuronal differentiation and maturation—at 1, 3, and 5 days post progenitor expansion (Fig. S2). Following the model described by Dotti et al. (1988), we analysed neuronal morphometry using micrographs obtained from cultures immunolabelled with βIII-Tubulin. Neuronal morphology was classified as follows: monopolar, exhibiting a single neuritic process; bipolar, exhibiting two processes; stage 2, exhibiting more than two processes of similar length; and stage 3, when one process was at least twice as long as the others. As shown in Fig. 3A, neurons from both genotypes (PS1^control^ and PS1^M146L^) differentiated/matured at a similar rate. At 1 DIV, the proportion of stage 3 neurons was lower than that of the other types, whereas at 3 and 5 DIV it became higher, regardless of genotype. Moreover, we did not notice any difference in cellular density between PS1^M146L^- and PS1^control^-iNs cultures during neuronal differentiation. However, when analysing neurite outgrowth, we observed differences in the total length of neuritic processes at 3 DIV. As shown in Fig. 3B, neurons derived from patients exhibited significantly increased neurite outgrowth at 3 DIV compared to the control group. Quantification was performed using the same micrographs previously analysed, employing ImageJ software. As a readout of excitability, we measured the Ca^2+^ uptake ability evoked by KCl stimulation in control and PS1 M146L iNs. As depicted in Fig. 3C-D, 14 DIV control iNs showed a consistent increase in cytosolic Ca2+ levels immediately after depolarisation. Nevertheless, PS1 ^M146L^ iNs failed to evoke a consistent response. As indicted in the Fig. 3D, just a few iNs displayed a mild increment in Ca^2+^ levels, whilst most of them remained unresponsive, suggesting a failure in consolidating their excitability capacity at this time point.

**Figure 3.**
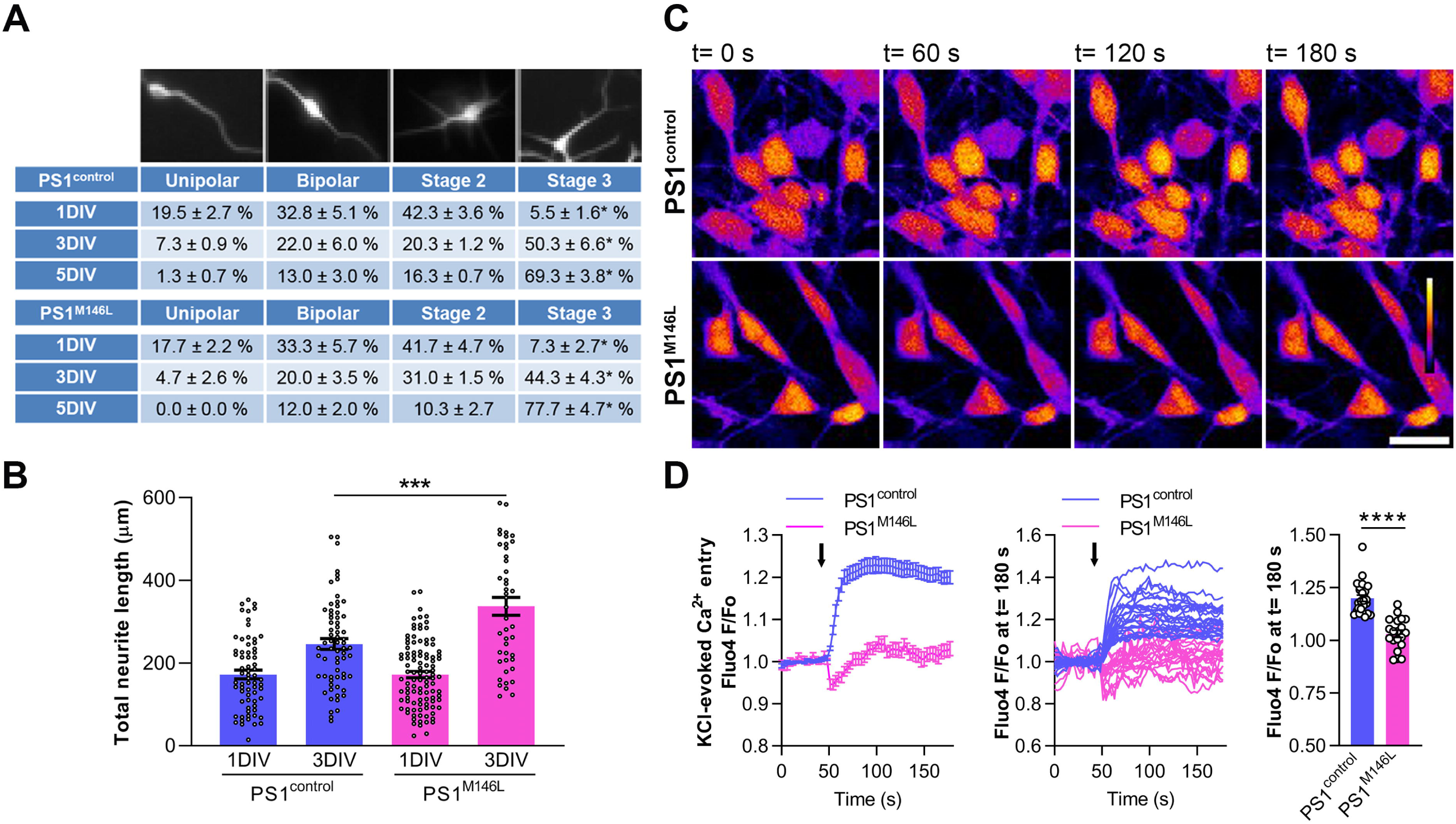
Morphological and physiological characterization of PS1^control^ and PS1^M146L^ iNs. **(A)** Morphological analysis of early PS1^control^ and PS1^M146L^ iNs in vitro. Percentage of unipolar, bipolar and multipolar stages of differentiation at 1, 3, and 5 DIV (data is shown as mean ± SEM of 3-4 biological replicates). Two-way ANOVA revealed no significant interaction between genotype and neuronal morphology at any time point (1 DIV: F=0.58, p=n.s.; 3 DIV: F=1.77, p=n.s.; 5 DIV: F=2.51, p=n.s.), indicating that neurons from both genotypes differentiated/matured at a comparable rate. In contrast, the main effect of morphology was significant across all time points (1 DIV: F=33.8, p<0.0001; 3 DIV: F=38.7, p<0.0001; 5 DIV: F=299.9, p<0.0001). Post-hoc Dunnett’s tests showed that at 1 DIV, the proportion of stage 3 neurons was significantly lower compared with other morphological stages (*p<0.05 for all comparisons), whereas at 3 and 5 DIV, stage 3 neurons were significantly more abundant than the other types (*p<0.05 for all comparisons). **(B)** Quantification of total neuritic length of PS1^control^ and PS1^M146L^ iNs. Bars and error bars show the mean ± SEM with individual values superimposed (t=3.63, ***p=0.0005, Welch’s correction). **(C)** Representative images of a recording field with PS1^control^ and PS1^M146L^ neurons at t=0, 60, 120 and 180 s. Scale bar: 20 µm. **(D) Left and middle panels.** Fluo4 time-lapse showing Ca^2+^ influx evoked by 120 mM KCl in PS1^control^ and PS1^M146L^ iNs at 14 DIV (left panel: average curve ± SEM; middle panel: individual values). The arrows indicate the time of KCl stimulation; **Right panel.** Statistical comparison of F/Fo ratios at t=180 s between PS1^control^ and PS1^M146L^ iNs. Bars and error bars show the mean ± SEM with individual values superimposed (t=7.4, ****p<0.0001).

The observed phenotype aligns with previous reports linking this genetic variant to defective synaptic transmission and could underlie the electrophysiological deficits associated with the disease (Wang et al., 2009). These findings further support the pathogenicity of the mutation and its potential role in disrupting neuronal network activity.

### Mitochondrial Ca^2+^ levels ([Ca^2+^]m) are significantly lower in PS1^M146L^ iNs under stressful conditions

Thapsigargin induces Ca^2+^ leak from the ER by inhibiting SERCA, which is potentially toxic for the cell. Consequently, mitochondria are critical for handling this response by uptaking Ca^2+^ into their inner space, mostly through the mitochondrial calcium uniporter 1 (MCU-1) (Márta et al., 2021). Thus, to evaluate the buffering capacity of mitochondria in PS1^control^ and PS1^M146L^ backgrounds, we used the mitochondrial-specific Ca^2+^-sensitive dye Rhod2-AM (Fernandez-Sanz et al., 2019; Marmolejo-Garza et al., 2023). Live neurons were imaged using a confocal time-lapse to record [Ca^2+^]_m_ challenged by 1 µM thapsigargin to promote cytosolic Ca^2+^ accumulation and thereby testing mitochondrial uptake. Our results suggest that PS1^M146L^ neurons display a significant reduction in their [Ca^2+^]_m_ levels, particularly at the equilibrium phase of the recording (Fig. 4A), suggesting failures in their [Ca^2+^]_m_ buffering capacity. We also assessed [Ca^2+^]_m_ levels in the presence of the MCU1 inhibitor (i11) in control (Fig. 4C) and PS1^M146L^ (Fig. 4E) neurons. For both control and PS1^M146L^ contexts, treatment with 50 µM i11 blocked the rise in [Ca^2+^]_m_ triggered by thapsigargin, reinforcing the notion that the MCU-1 is the main gate for mitochondrial Ca^2+^ uptake.

**Figure 4.**
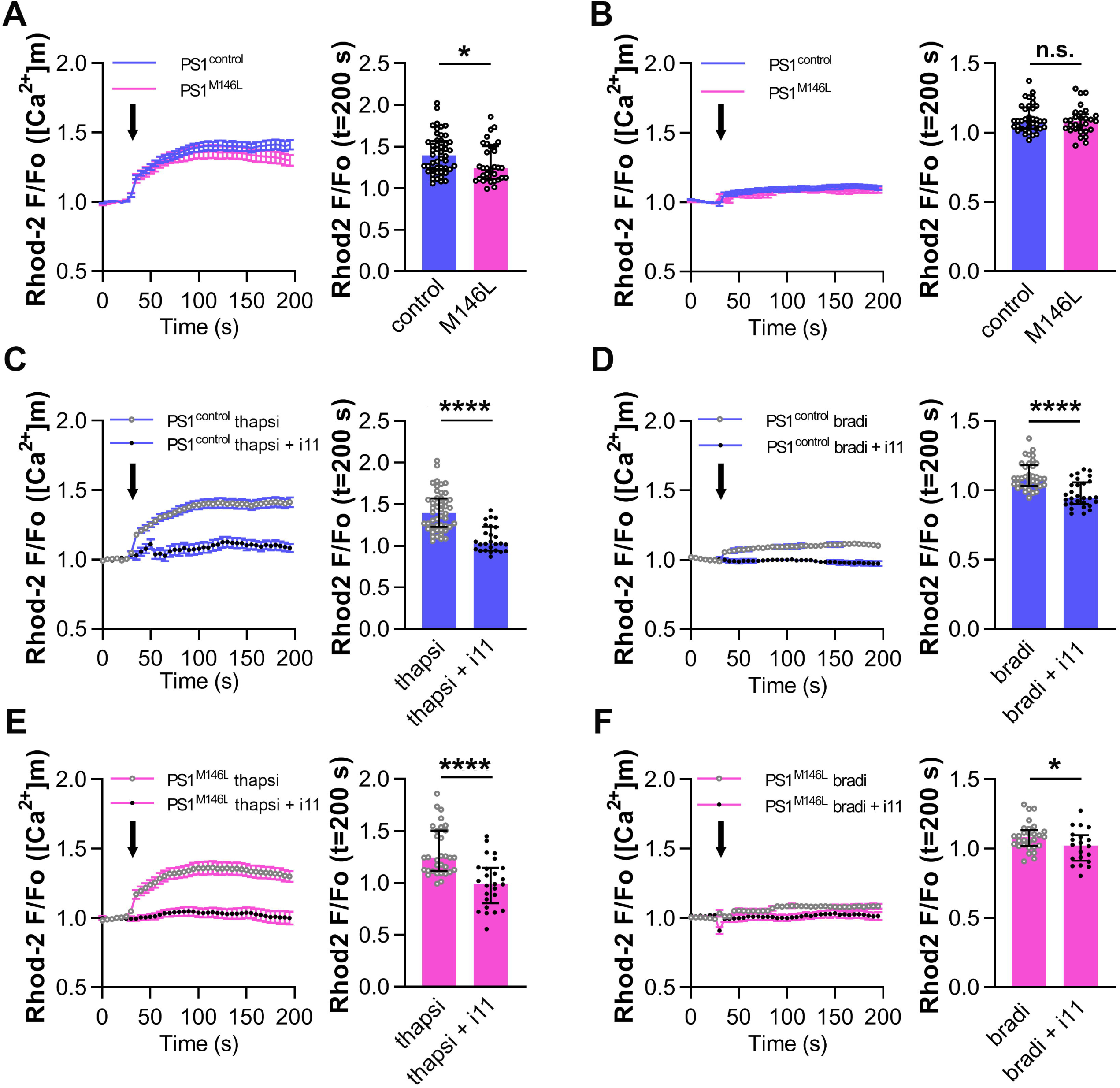
Evaluation of mitochondrial Ca^2+^ ([Ca²⁺]m) levels in PS1^control^ and PS1^M146L^ iNs. Seven DIV PS1^control^ and PS1^M146L^ iNs were stimulated with 1 µM thapsigargin (A, C, E) or 200 nM bradykinin (B, D, F). [Ca²⁺]m levels were recorded using the mitochondrial Ca^2+^ reporter Rhod2-AM. **(A)** Comparison of [Ca²⁺]m between PS1^control^ and PS1^M146L^ iNs challenged by thapsigargin (U=536, *p=0.0222, Mann-Whitney test). **(C)** [Ca²⁺]m levels in PS1^control^ iNs in the presence or absence of the MCU inhibitor i11, challenged by thapsigargin (U=144, ****p<0.0001, Mann-Whitney test). **(E)** [Ca²⁺]m levels in PS1^M146L^ iNs in the presence or absence of the MCU inhibitor i11, challenged by thapsigargin (U=144.5, ****p<0.0001, Mann-Whitney test). **(B)** Comparison of [Ca²⁺]m between PS1^control^ and PS1^M146L^ iNs challenged by bradykinin (U=499, p=n.s., Mann-Whitney test). **(D)** [Ca²⁺]m levels in PS1^control^ iNs in the presence or absence of the MCU inhibitor i11, challenged by bradykinin (U=200.5, ****p<0.0001, Mann-Whitney test). **(F)** [Ca²⁺]m levels in PS1^M146L^ iNs in the presence or absence of the MCU inhibitor i11, challenged by bradykinin (U=192.5, *p=0.0472). In the right panels, the data are presented as mean ± SEM. In the left panels, the bars and error bars show the median with the interquartile range (individual values superimposed). N=3 biological replicates.

In addition, we evaluated the contribution of IP3R to [Ca^2+^]_m_, as it is well documented that IP3R enables the flow of Ca^2+^ from the ER to mitochondria (Filadi et al., 2016; Jaconi et al., 2000; Zampese et al., 2011). Accordingly, PS1^control^ and PS1^M146L^ neurons were challenged by the IP3R agonist bradykinin (200 nM), triggering the activation of G protein-coupled receptors (B1R/B2R), phospholipase C (PLC), generation of IP3 and ER Ca²⁺ release through IP3R. However, PS1^control^ and PS1^M146L^ neurons responded to a similar extent (Fig. 4B, 4D and 4F), suggesting that IP3R-dependent Ca²⁺ mitochondrial uptake (MCU) functions normally in this specific context.

Taken together, our findings indicate that the PS1^M146L^ mutation compromises the ability of mitochondria to buffer Ca^2+^ dysregulations under stressful conditions.

### The PS1^M146L^ mutation leads to abnormal mitochondrial function and structure

Neurons with altered metabolism may have a reduced ability to maintain ionic gradients (Attwell and Laughlin, 2001). To assess mitochondrial function, we measured respiration in PS1^control^ and PS1^M146L^ iNs at 21DIV using a Seahorse XFp extracellular flux analyser (Agilent), which records oxygen consumption rate (OCR) in intact cells. By applying specific ETC modulators, we quantified key bioenergetic parameters. Oligomycin, an ATP synthase inhibitor, was employed to determine basal respiration associated to the production of ATP; FCCP, a protonophore that collapses the mitochondrial membrane potential, revealed maximal respiratory capacity; finally, rotenone and antimycin A (complex I and III inhibitors, respectively) were added to evaluate non-mitochondrial respiration (Fig. 5A). PS1^M146L^ iNs showed significantly higher values than PS1^control^ across all parameters except proton leak (Fig. 5B). Proton leak reflects the return of protons across the inner mitochondrial membrane without ATP synthesis, dissipating the gradient as heat. Comparable leak levels indicate that energy dissipation is unchanged, but dissimilarities in other parameters suggest a distinct metabolic state. Elevated basal respiration in PS1^M146L^ cells points to a higher energy demand at rest, consistent with increased mitochondrial ATP production to sustain cellular requirements. The stronger response to FCCP indicates that substrate oxidation and respiratory capacity are greater in PS1^M146L^ than in PS1^control^. In addition, PS1^M146L^ cells displayed higher non-mitochondrial respiration after rotenone/antimycin A treatment, which may reflect increased NADPH oxidase (NOX family) activity. NOX enzymes generate superoxide (O₂⁻) and H₂O₂ by transferring electrons from NADPH to O₂, processes relevant to both neuronal physiology and pathology (Bedard et al., 2007).

**Figure 5.**
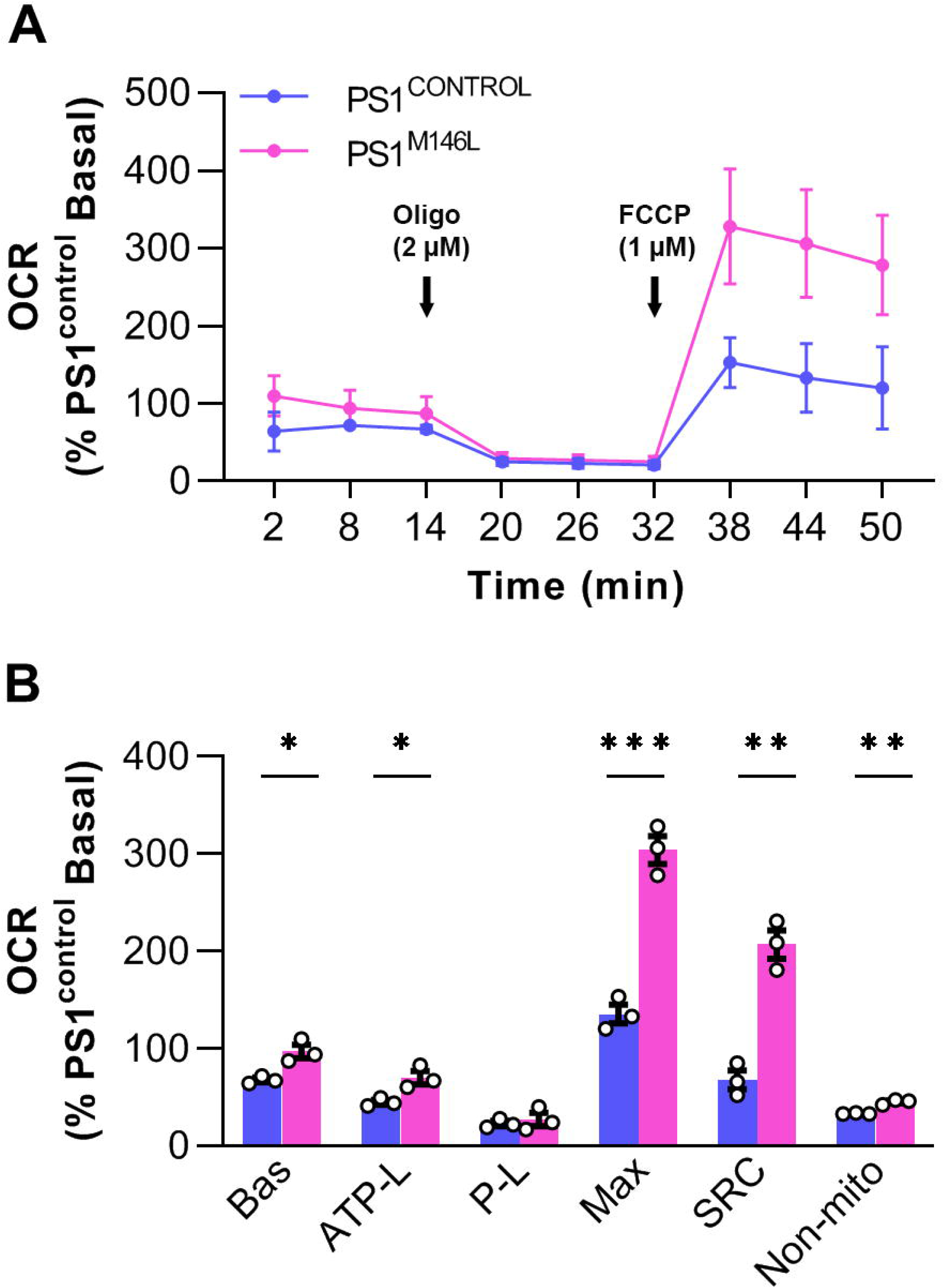
Mitochondrial functionality of PS1^M146L^ iNs. **(A)** Representative profile of the oxygen consumption rate (OCR) of iNs from PS1^control^ and PS1^M146L^. OCR was determined in the presence of 25 mM glucose plus 1 mM pyruvate and 2 mM of glutamine as substrates. Graph shows basal OCR, OCR after ATP synthase inhibition by 2 µM oligomycin and maximum OCR after the addition of 1 µM FCCP. Data are shown as mean ± SEM. **(B)** Bars show the oxygen consumption rate (OCR) of iNs from PS1^control^ and PS1^M146L^ expressed as percentage of PS1^control^ basal value. Bars and error bars show the mean ± SEM with individual values superimposed (Bas, t=4.08, p=*0.015; ATP-L, t=3.52, p=*0.024; P-L, t=0.55, p=n.s.; Max, t=9.71, p=***0.0006; SRC, t=8.03, **p=0.0013; Non-mito, t=7.46, **p=0.0017).

Together, these findings indicate that PS1^M146L^ iNs may generate more mitochondrial ATP than their non-mutated counterparts. The higher maximal respiration could reflect either enhanced mitochondrial function or an expansion of the mitochondrial network. To address this, we evaluated whether the increased respiratory parameters were simply due to greater cell size or mitochondrial mass by calculating coupling efficiency, spare respiratory capacity (SRC), and the respiratory control ratio (RCR) (Table 1). These indices are expressed as ratios of respiration under different conditions, providing internally normalised measures that are independent of cell content or protein mass (Brand and Nicholls, 2011).

**Table 1.** Mitochondrial respiratory ratios.

| Respiratory Ratios | HCS | PS1<sup>M146L</sup> |
| --- | --- | --- |
| Coupling efficiency | 0.659 ± 0.036 | 0.721 ± 0.069 |
| Spare Respiratory Capacity (SRC) | 2.000 ± 0.141 | 3.132 ± 0.150** |
| Respiratory control ratio (RCR) | 5.860 ± 0.414 | 11.236 ± 0.537** |

Coupling efficiency, which refers to the proportion of basal respiration used to ATP synthesis, did not differ between PS1^M146L^ and PS1^control^ iNs (Table 1), indicating that electron transport was similarly coupled to ADP phosphorylation in both groups. In contrast, SRC (maximal - basal respiration) was higher in PS1^M146L^ cells (Table 1), consistent with increased ATP demand. Likewise, the RCR (maximal respiration relative to oligomycin-resistant respiration) was elevated in PS1^M146L^ compared with PS1^control^ (Table 1), suggesting enhanced mitochondrial respiratory function.

To test whether the increased mitochondrial activity in PS1^M146L^ iNs was due to greater mitochondrial content, we measured TOM20 protein levels (Fig. S3A, B) and mtDNA-to-nDNA ratios (Fig. S3C). Neither parameter differed between PS1^M146L^ and PS1control cells. In addition, the higher ATP production observed in PS1^M146L^ iNs was not explained by changes in ATP synthase (ATPase) transcript levels (Fig. S3D).

We next analysed mitochondrial network by confocal microscopy after labelling cells with MitoTracker Green FM. Short, rounded mitochondria were predominantly displayed by PS1^control^ iNs, whereas elongated, tubular structures were shown by PS1M146L neurons (Fig. 6A). Quantification confirmed longer mitochondria in PS1^M146L^ compared with PS1^control^ (Fig. 6B). Consistently, mitofusin 1 (Mfn1) expression was higher in PS1^M146L^ cells (Fig. 6C-D), supporting the observed mitochondrial elongation.

**Figure 6.**
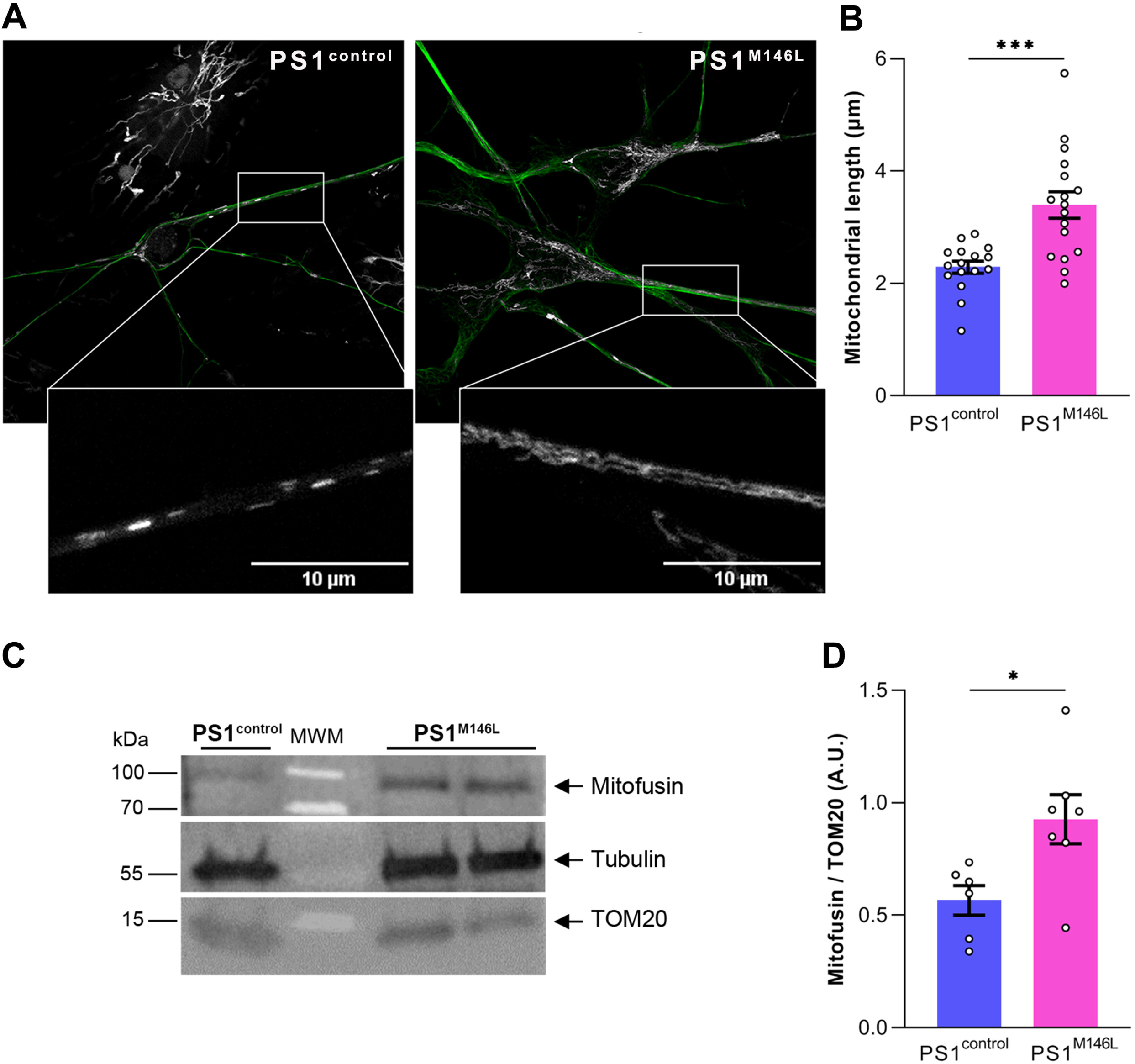
PS1^M146L^ iNs present abnormal morphology. **(A)** Left panel. Representative images of living cells stained with MitoTracker Green FM and analysed by confocal microscopy (×630). PS1^control^ (n=16) and PS1^M146L^ (n=17) photomicrographs were assessed. Magnified images of the boxed regions are shown below. Scale bars, 10 µm. **B)** Bars and error bars represent the mean ± SEM (individual values superimposed) of mitochondrial length from the images analysed in panel A (t=4.28, ***p=0.0003, Welch’s correction). **(C)** Representative Western blots for mitochondrial fusion protein Mitofusin (MFN), TOM20 and Tubulin (n = 3-5 culture dishes per group). **D)** Bars and error bars represent the mean ± SEM (individual values superimposed) of Mitofusin normalized by the total amount of mitochondria (TOM20) (t=2.72, *p=0.02).

To further explore the link between Ca2+ homeostasis and oxidative stress, we measured reactive oxygen species (ROS) in live, neuron-like cells using the fluorescent probe dichlorodihydrofluorescein diacetate (DCF-DA) (Fig. 7A). As shown in Fig. 7B, ROS levels were markedly higher in PS1^M146L^ iNs compared with PS1^control^, consistent with the observed enhanced mitochondrial activity.

**Figure 7.**
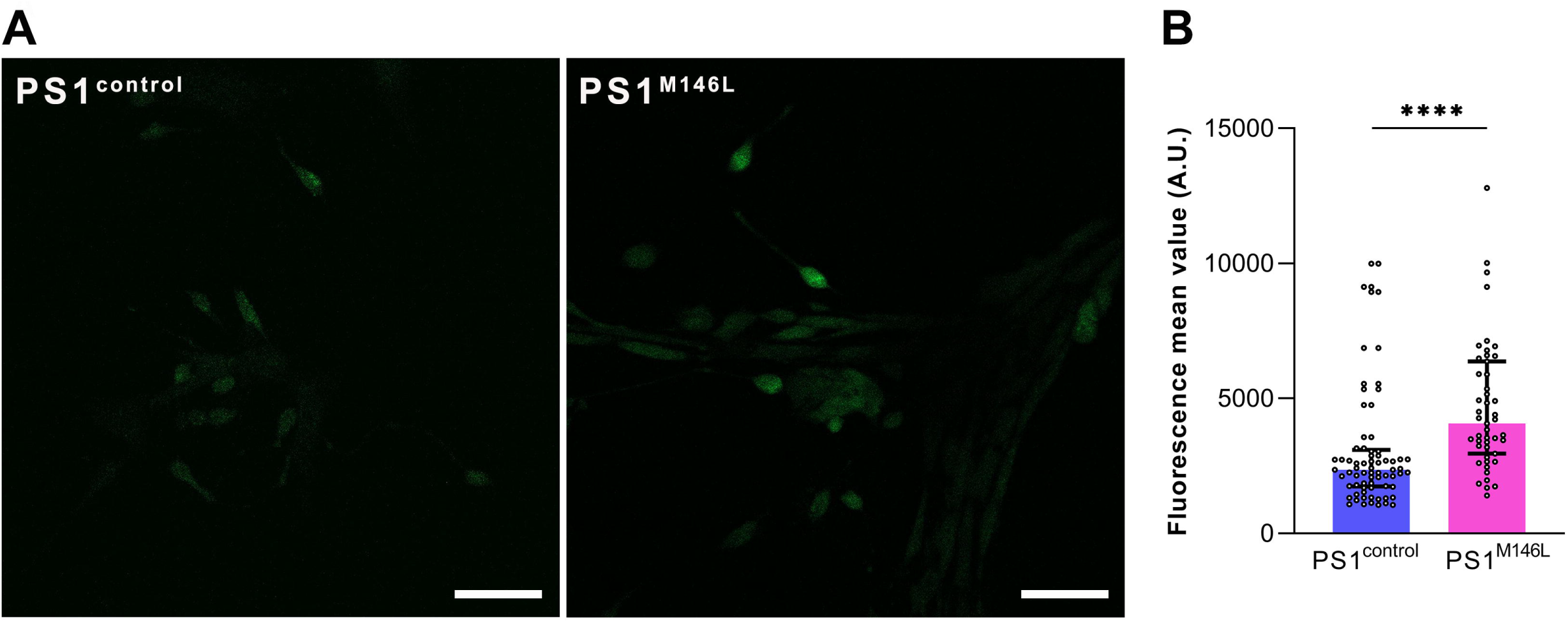
Reactive oxygen species (ROS) determined by DCF in iNs. **(A)** Representative microphotographs showing DCF fluorescence intensity of total ROS level in PS1^control^ and PS1^M146L^ neurons. Scale bars, 50 µm. **(B)** Quantitative intracellular DCF fluorescence intensity per neuron. Bars and error bars show the median and the interquartile range with individual values superimposed (U=857, ****p<0.0001, Mann-Whitney test).

## Discussion

Current insights into FAD mechanisms largely rely on animal models, which do not fully recapitulate human disease due to fundamental differences between rodent and human brains (Oberheim et al., 2009). Using iNs derived from a PS1^M146L^ FAD carrier, we show that this mutation affects Aβ42 production, calcium dynamics, and mitochondrial activity and morphology, highlighting the early dysregulation of critical intracellular signalling pathways in neurons from at-risk individuals.

It is notable that iNs were not stressed at any point in order to reveal these abnormalities; they were undistinguishable under usual culture conditions. We observed no significant differences in cell density or overall morphology between PS1^M146L^ and PS1^control^ neurons. The only exception was a faster initial neurite outgrowth in PS1^M146L^ iNs at 3 DIV. These observations are in agreement with those reported by Yang et al. (2017) at 28 DIV, where neural stem cell cultures from FAD NPCs displayed spontaneous differentiation with extensive neurite outgrowth, while cells from healthy NPCs retained typical progenitor morphology.

A hallmark of FAD neuropathology is the accumulation of Aβ peptides in the brain. In our study, PS1^M146L^ iNs did not show increased production of Aβ38 or Aβ40 but did exhibit elevated Aβ42 levels and a higher Aβ42/40 ratio. These results agree with earlier findings indicating increased generation of the aggregation-prone Aβ42 in both the brains and peripheral cells of mutation carriers (Scheuner et al., 1996). Consequently, PS1^M146L^ iNs may provide a valuable platform for testing anti-amyloid therapies aimed at modulating Aβ42 production, clearance, or toxicity in a human neuronal context.

In this study, using multiple live-cell imaging protocols, we evaluated cytosolic and mitochondrial Ca^2+^ dynamics in iNs from PS1^control^ and PS1^M146L^ donors. Calcium homeostasis is critical for neuronal physiology at many levels, ranging from differentiation to neurotransmission and brain connectivity (Angelova and Abramov, 2024; Doser and Hoerndli, 2021). Therefore, perturbations in Ca^2+^ signalling, especially when chronic, rapidly turn into cell dysfunction and pathology. Emerging evidence suggests a critical contribution of PS2 to preserve ER-mitochondria contacts and Ca^2+^ crosstalk between both organelles (Zampese et al., 2011). Particularly, mutations in PS2 lead to Ca^2+^ leak from ER by inhibiting SERCA in neuroblastoma cell lines (Filadi et al., 2016; Zampese et al., 2011). However, the contribution of PS1 mutations to Ca^2+^ dynamics in human neurons remains to be probed. Using the cytosolic Ca^2+^ reporter Fluo4, here we report that PS1^M146L^ iNs failed to evoke a depolarizing response, at least within the time in culture analyzed in this study. In this regard, several hypotheses may explain this behavior, including changes in membrane permeability and composition, voltage-dependent ion channels expression, and the maturational stage, among others. Moreover, using the Ca^2+^ reporter Rhod2, we detected that PS1^M146L^ mitochondria uptake significantly less Ca^2+^ after thapsigargin-dependent ER depletion mostly mediated by the MCU-1. This data suggests that this mutation impairs the capacity of mitochondria to buffer Ca^2+^ dysregulations under stressful conditions.

Physiologically, the synthesis of inositol 1,4,5-triphosphate (IP3) promotes Ca^2+^ leak from the IP3R at the ER surface to mitochondria (Jaconi et al., 2000). To test this hypothesis under the PS1^M146L^ context, we measured [Ca²⁺]m dynamics in control and mutant iNs challenged by bradykinin. Our data unveiled no differences between the genetic backgrounds, suggesting that PS1^M146L^ does not impact IP3-mediated [Ca²⁺]m uptake. Alternatively, it is also possible that the impact of the mutation is not evident because bradykinin activates compensatory mechanisms, such as store-operated calcium entry (SOCE) that could mask defects. In any case, Ca^2+^ signals evoked by bradykinin were mild in PS1 ^control^ and PS1 ^M146L^ iNs, suggesting IP3R-mediated signaling could not be fully consolidated within the timeframe of this study.

Neurons are highly metabolically active cells (Bélanger et al., 2011) and are particularly prone to oxidative stress due to their elevated oxygen consumption, limited antioxidant capacity, and the dense organization of brain tissue (Wilson et al., 2015). Ca^2+^ signalling is closely linked to reactive oxygen species (ROS) production, especially when dysregulated (Bórquez et al., 2016; Muñoz-Palma et al., 2025 preprint; Wilson et al., 2016, 2018). Previous in situ respirometry studies, including our own, have revealed alterations in energy metabolism in both human and animal models of Alzheimer’s disease (AD), with outcomes varying according to the specific FAD mutation and cellular model examined (Area-Gomez et al., 2012; Martino Adami et al., 2019). A detailed analysis of how the PS1^M146L^ mutation affects mitochondrial function and Ca^2+^ regulation in human neurons has not yet been reported. Here, we show that PS1^M146L^ enhances oxidative phosphorylation and modifies mitochondrial morphology in iNs. Our metabolic flux data are consistent with earlier reports of increased mitochondria ATP-linked respiration, and SRC in fibroblasts from PS1 mutation carriers (mean age 48.3 years) (Bell et al., 2018). By contrast, a recent study in H4 glioblastoma cells engineered to inducibly express either wild-type PS1 or five PS1 mutants (excluding M146L) reported only minor effects of PS1 mutations on mitochondrial bioenergetics (Han et al., 2021). The discrepancy with our findings likely reflects differences in the FAD mutation examined, the neuronal cell type studied, and the genetic manipulations performed, underscoring the value of studying human neurons to better understand disease-relevant mechanisms.

Our findings indicate that an elevated basal oxygen consumption rate, in conjunction with aberrant mitochondrial calcium homeostasis, unveils a paradoxical metabolic phenotype in PS1^M146L^ neurons. This observation lends support to the hypothesis that PS1^M146L^ neurons manifest compensatory mitochondrial hyperfunction. The results obtained suggest that PS1^M146L^ neurons undergo a loss of normal Ca²⁺ regulation, with the neurons compensating for this by increasing their respiratory activity to maintain ATP levels. The elevated oxygen consumption exhibited by mutant neurons is plausible as a maladaptive response. In this regard, the loss of normal Ca²⁺ signalling could trigger the activation of energetically costly alternative Ca²⁺ pathways (SOCE), forcing mitochondria to expand their capacity to maintain homeostasis. In this sense, mutant neurons could develop a hyperactive metabolic phenotype as an early compensation for cellular stress, before the energy collapse that is typical of advanced stages of FAD pathology. In agreement with these results, functional experiments performed on isolated mitochondria from the hippocampus of two APP knock-in AD mouse models, confirmed that hypermetabolism of the mitochondria appears before impaired autophagy and synaptic disorganisation (Naia et al. 2023).

Intriguingly, FAD fibroblasts holding PS1 mutations exhibit abnormally high resting levels of ROS, leading to stress-induced premature senescence (Ma et al., 2014). In this work, we determined cytosolic ROS levels using the fluorescent probe DCF, unveiling a notable increase in PS1^M146L^ iNs, which is strongly correlated with Ca^2+^ perturbations observed in these neurons. Physiological levels of ROS are critical to maintain the activity of Ca^2+^ channels, including IP3R, RyR, NMDAR and L-type channels (Muñoz-Palma et al., 2025 preprint; Wilson et al., 2016). Thus, our data suggest that mutant neurons exhibit a global dysregulation of critical intracellular messengers, such as Ca^2+^ and ROS, ultimately affecting neuronal functions. Consequently, the enhancement in spare respiratory capacity may be regarded as an early protective response before the onset of neurodegeneration. However, this metabolic alternative is associated with an elevated risk of progressive substrate depletion and ROS accumulation, which would result in mitochondrial dysfunction, a hallmark of advanced stages of the disease.

In this study, we observed increased mitochondrial length and elevated expression of the mitochondrial fusion protein mitofusin 1 (Mfn1) in iNs from PS1^M146L^ carriers compared with PS1^control^ cells. The presence of elongated mitochondria is a classic hallmark of altered mitophagy. In this regard, our results are in agreements with a previous report showing mitophagy failure in iPSC-derived neurons of PS1 A246E mutation (Martín-Maestro et al., 2017). Mitochondrial fusion is a complex process involving the merging of four membranes. In mammalian cells, key components include mitofusin 1 (MFN1) and mitofusin 2 (MFN2) for outer membrane fusion, OPA1 for inner membrane fusion, and various accessory and regulatory proteins. Fusion is essential for maintaining a mitochondrial population with a complete set of nuclear- and mitochondrial-encoded gene products and typically increases under conditions that demand higher mitochondrial ATP production. In the context of Alzheimer’s disease, fusion allows damaged mitochondria to merge with functional ones, diluting and reorganising the mitochondrial matrix. Highly interconnected mitochondrial networks suggest that fusion contributes to energy management, with mitochondrial morphology reflecting cellular needs (Westermann, 2010). Altered mitochondrial dynamics are also a hallmark of cellular senescence, where mitochondria often appear elongated, enlarged, and hyperfused. This hyperfused state may protect against apoptosis. While senescence has been documented in post-mitotic neurons (Jurk et al., 2012; Piechota et al., 2016), its occurrence in PS1^M146L^ iNs has not yet been investigated.

Collectively, our data show that post-mitotic iNs from a PS1^M146L^ carrier show a profile compatible with some of the senescence makers, shedding light on inconsistent results due to different neuronal culture conditions and measurements.

Furthermore, our results indicate that in early mutant neurons, an imbalance of Ca²⁺ signalling induces an energy-intensive metabolic response to maintain cellular homeostasis. These findings further support the idea that changes in spatiotemporal dynamics of Ca²⁺ should be considered an early event in the pathophysiology of AD, and a crucial target for early interventions.

## Materials and Methods

### Enrolment of participants

We performed detailed neuropsychological tests and skin biopsies for iPSC generation from each participant. The Institutional Review Board (IRB) of Fundación Instituto Leloir (#00007572) approved the protocol, in accordance with its regulations. Dermal punch biopsies were taken following informed consent at the Centre of Neuropsychiatry and Neurology of Behaviour (School of Medicine of the University of Buenos Aires, Argentina).

### Clinical history of fibroblast donors

Subject #1: Carrier of PS1^M146L.^. Sex, female; ApoE genotype, 3/3; Diagnosis, FAD carrier; Age at sampling: 35 years. Member of a four-generation Argentine pedigree (ARI) in which early-onset FAD segregates as an autosomal dominant trait (Morelli et al., 1998). At the age of 35, she had normal neuropsychological examination results and was considered an asymptomatic family member.

Subject #2. Non-carrier of PS1^M146L^. Sex, female; ApoE genotype, 3/3; Diagnosis, healthy control; Age at sampling: 39 years.

### Cell culture and neural differentiation

iPSC clones (two clones for each subject) derived from PS1^M146L^ and PS1^control^ fibroblasts were obtained from PLACEMA (Fundación Instituto Leloir) and cultured on irradiated mouse embryonic fibroblasts (MEFi) according to standard methods. iPSCs were differentiated into neurons as described in Zhang and Zhang (2010), with minor modifications. Human iPSCs were dissociated with trypsin (Thermo Fisher Scientific) for 2 min at 37 °C / 5% CO_2_. Embryoid bodies (EBs) were generated by transferring dissociated iPSCs to a non-adherent U-bottom 96-well plate (15,000 cells per well, 150 μL of EB medium (DMEM/F12 supplemented with 0.1 mM 2-ME, 0.1 mM NEAA, 2 mM Glutamax, and 20% (vol/vol) KSR) plus 4 ng/mL human FGF2 and 50 μM Y27632 compound (Calbiochem), allowing reaggregation of cells within 24 h. Cell aggregates were transferred onto a 24-well plate and maintained in suspension for 5 days in EB medium, and then, 2 extra days in neural induction medium (NIM: DMEM/F12, 1% N2 supplement (Thermo Fisher Scientific), 2 μg/ mL heparin (Sigma-Aldrich) and 1% penicillin/streptomycin). For the generation of neural precursor cells (NPCs) as neural rosettes, EBs were plated onto Geltrex-coated dishes (Invitrogen), in NIM (d7–14). Neural rosettes were harvested mechanically and replated on non-treated culture dishes in NIM for neurosphere formation, and maintained for an additional 15 days. From day 28, neural progenitors were differentiated into neurons on a Poly-L-Lysine/Geltrex substrate in neuronal differentiation medium (NDM) consisting of Neurobasal medium, B27 supplement, cAMP (1 μM), BDNF, GDNF, IGF1 (10 ng/mL) and Ascorbic Acid.

### Characterisation of iNs by immunocytochemistry

Fixation of cells was performed using 4% paraformaldehyde and 4% sucrose in PBS at RT for 20 min. Cells were then permeabilised with 0.2 % PBS-Triton and blocked with 5% bovine serum albumin in 0.1% PBS-Tween during 1 h. Next, cells were incubated with primary antibodies at RT for 1 h. The following primary antibodies were used, either individually or in combination: rabbit anti-SOX2 (1:100, Abcam, ab137285), mouse anti-nestin (1:500, Abcam, ab22035), mouse anti-βIII tubulin (1:1000, Abcam, ab78078), and chicken anti-MAP2 (1:500, BioLegend, 822501). Subsequently, cells were washed and incubated with secondary antibodies at RT for 1 h. The following second antibodies were used: anti-rabbit IgG (1:1000, Alexa Fluor 633, Life Technologies, A21070), anti-mouse IgG (1:1000, Alexa Fluor 488 and 546; Life Technologies, A11029 and A11004), and anti-chicken IgY (1:1000, Alexa Fluor 488, Invitrogen, A11039). Zeiss LSM 800 confocal microscope was employed to acquire images at the “Centro de Micro y Nanoscopía de Córdoba” (CEMINCO-CONICET-UNC). Total neurite length, calculated as the sum of all branches in each individual neuron labelled with anti-βIII tubulin, was quantified using ImageJ (NIH, USA). Quantification of the percentage of positive cells for each antibody over DAPI positive cells were performed using an InCell Analyzer 2500 HS (GE Healthcare) and analysed with IN Carta Image Analysis software.

### Live-cell cytosolic Ca^2+^ imaging

iNs were plated into 25 mm-diameter glass coverslips (pre-coated with Poly-L-Lysine and Geltrex). The cells were maintained in Neurobasal medium (supplemented with B27 and Glutamax) until the time-lapse assay. Cells were incubated with 5 µM Fluo4-AM (Life Technologies, USA) for 20 min at 37 °C (cell incubator) in Tyrode’s buffer (TB: 110 mM NaCl, 2.5 mM KCl, 2 mM CaCl_2_, 2 mM MgCl_2_, 25 mM Hepes, 30 mM glucose). Then, TB was discarded and replaced by Neurobasal with supplements for 30 min at 37 °C (Mir et al., 2020; Wilson et al., 2016). Later, the coverslips were transferred to a cell perfusion chamber holding 500 μL TB. Imaging was done in a Zeiss LSM 800 confocal microscope (40x, NA=1.4, pinhole=1 AU), using the time-lapse configuration to obtain 512 x 512 px frames every 2 s. Samples were excited with a laser λexc=488 nm, and emission was collected within the λem=505-530 nm range. A 2x KCl (or 2x glycine) solution was added to the cell chamber (500 μL, 240 mM KCl; 400 μM glycine), previously filled with 500 μL TB, after the first 30 s of recording (baseline). To estimate Fluo4 fluorescence as a function of time [F(t)], time-lapse files were analysed using the Time Series Analyzer plugin in Fiji-ImageJ, defining a circular ROI within the soma of each cell. The Fo value corresponds to the average F(t) of the baseline. The ratio F(t)/Fo represents the fold change of Fluo4 intensity over time.

### Live-cell mitochondrial Ca^2+^ imaging

Cells were cultured as indicated for the cytosolic Ca^2+^ recordings until the time of the assay. Then, iNs were incubated with 5 μM Rhod-2 AM (LifeTechnologies, USA) in mitochondrial Ca²⁺ -free imaging buffer (MIB: 156 mM NaCl, 3 mM KCl, 2 mM MgSO_4_, 1.25 mM K_2_HPO_4_, 2 mM CaCl_2_, 10 mM Hepes, 10 mM D-glucose, 1 mM EGTA) for 30 min at RT (on the bench, protected from light exposure), followed by 20 min at 37 °C (cell incubator) (Fernandez-Sanz et al., 2019; Marmolejo-Garza et al., 2023). When needed, cells were treated with 50 μM MCU inhibitor (i11) from this time, and for at least 30 min before the time-lapse assay (37 °C, cell incubator). Later, the coverslips were transferred to a cell perfusion chamber holding 500 μL MIB. Imaging was done in a Zeiss LSM 800 confocal microscope (40x, NA=1.4, pinhole=1 AU), using the time-lapse configuration to obtain 512 x 512 px frames every 5 s. Samples were excited with a laser λexc=561 nm, and emission was collected within λem=570-600 nm range. After the baseline (first 30 s), 500 μL of 2x stimuli were added (2 μM thapsigargin, 200 nM bradykinin). Time-lapse files were analysed using the Time Series Analyzer plugin in Fiji-ImageJ, as indicated for cytosolic Ca^2+^ recordings.

### Measurement of ROS by fluorescence microscopy

Neural progenitors were differentiated on coverslips for 7 days as described above. 7 DIV neurons were incubated (10 min at 37 °C) with 1 μM Dichorofluorescein diacetate (DHF, Sigma D6883). Then, neurons were washed to remove dead cells and debris. After loading, the coverslip was mounted in a plastic chamber and covered with 900 µL PBS/10mM Glucose/10mM Hepes. Photographs were taken in a Zeiss LSM 800 confocal microscope at 20x. Image analysis was performed offline using ImageJ software. For each neuron, a region of interest (ROI) was defined within cell soma and the average fluorescence intensity within each ROI was measured.

### Cellular bioenergetics

Oxygen consumption rate (OCR) was assessed with a Seahorse XFp extracellular flux Analyzer (Agilent). Culture medium was replaced with unbuffered DMEM (pH 7.4, and supplemented with 25 mM D-glucose, 1 mM sodium pyruvate, and 2 mM glutamine), prior to the assay. Then, cells were incubated for 1 h at 37 °C in the absence of CO₂. Basal OCR was recorded and subsequently monitored following sequential injections of oligomycin (2 μM), FCCP (1 μM; carbonyl cyanide 4-(trifluoromethoxy)phenylhydrazone), and antimycin A/rotenone (0.5 μM, AA/R). Non-mitochondrial respiration (AA/R-resistant) was subtracted from all OCR values. After the assay, the cells were fixed and stained with DAPI (1:500 in PBS) for counting. The number of nuclei was then quantified using an InCell Analyzer (GE Healthcare). OCR values are presented normalised to the number of nuclei.

### Mitochondrial morphology

To characterise the mitochondria, 15-DIV iNs plated on coverslips were incubated with 150nM MitoTracker Red (Invitrogen, M7512) in culture medium for 30 minutes at 37°C. The cells were then fixed, washed and immunostained with a mouse anti-tyrosine-tubulin antibody (diluted 1:1000; Sigma, T9028), followed by an Alexa Fluor 488-conjugated anti-mouse IgG secondary antibody (diluted 1:1000). Images were acquired using a Zeiss LSM 800 confocal microscope at CEMINCO-CONICET-UNC. Mitochondrial length was determined in the periphery region of the cells since in the perinuclear region, the high density of mitochondria prevented determining its size (Martinez et al., 2019). Sixteen photomicrographs of iNs of PS1^control^ and 17 of PS1^M146L^ mutation were analysed. In each photo, a mean value of mitochondrial length was determined, and the means were compared between genotypes. A total of 1,095 mitochondria (PS1^control^) and 696 individual mitochondria objects (PS1^M146L^) were segmented and analysed. Image processing and analysis was carried out using ImageJ 1.53t. Mitochondrial length was evaluated by measuring the Feret’s diameter of the segmented particles from the Max-Z project of the MitoTracker channel. Masks of whole βIII-tubulin (Tuj1)-positive cells were obtained from the maximum intensity projection by subtracting 70 intensity units from the background, applying first a median (radius 5 pixels), then an unsharp (radius=3 pixels, weight= 0.7) filter and finally thresholding using the Triangle method. Only particles with an area bigger than 600 px^2^ were considered. Somas of βIII-tubulin (Tuj1)-positive cells were selected manually. The final masks of the cell periphery resulted from the exclusive union of the whole cell masks and their respective soma masks. Total mitochondria were segmented using the intersection of two partial masks of the MitoTracker channel. The first one was obtained from a maximum intensity Z projection by normalising contrast to 0.3% pixel saturation and then applying an unsharp filter (radius= 100 pixels, weight=0.2) and thresholding with Li’s method. The second mask was obtained from the average intensity Z projection, to which a Gaussian blurred copy (sigma=15) was subtracted, and using the Max Entropy method for thresholding.

### Expression of Aβ isoforms in conditioned supernatants

MSD® V-PLEX PLUS Aβ Peptide Panel 1 kit was used to measure the levels of human Aβ38, Aβ40, and Aβ42 peptides secreted by iNs as previously reported (Galeano et al., 2023). Conditioned supernatants from four PS1^control^ and eight PS1^M146L^ samples were applied to a MULTI-SPOT® microplate that was pre-coated with and antibody raised against the C-termini of Aβ38, Aβ40, and Aβ42. SULFO-TAG™-labelled 6E10 monoclonal antibody was used for detection. Electrochemically stimulated light emission was detected with the MSD QuickPlex SQ120, and subsequent data analysis was performed using MSD Workbench 4.0 software.

### Assessment of mitochondrial (mt)DNA/nuclear (n)DNA ratio

Total DNA was extracted from iNs using the Illustra TriplePrep kit (GE Healthcare). DNA concentration was measured at A260 nm, and purity was evaluated via the A260/A280 ratio using a NanoDrop (Thermo Fisher Scientific). The following primers were used: mitochondrial encoded gene S1-ETC, 5-CAAACCTACGCCAAAATCCA-3 (forward) and 5-GAAATGAATGAGCCTACAGA-3 (reverse); nuclear 18S rRNA, 5-ACGGACCAGAGCGAAAGCA-3 (forward) and 5-GACATCTAAGGGCATCACAGAC-3 (reverse) (Sheng et al., 2012). Using FastStart Universal SYBR Green Master (ROX) and 10 μM primers, genomic DNA (50 ng) was amplified by qPCR in a LightCycler 480 (Roche). mtDNA and nDNA were amplified in separate reactions, and the mtDNA/nDNA ratio was calculated as 2^-(CtETC-Ct18S)^, and expressed as fold change relative to PS1^control^.

### Gene expression analysis of ATPase transcript levels

Illustra TriplePrep kit (GE Healthcare) was employed to extract total RNA from iNs. To determine the concentration and purity of RNA, absorbance at 260 nm and the ratio 260/280 were measured in a NanoDrop (Thermo Fisher Scientific). Reverse transcription was conducted in 20 μL containing: 200 ng of random primers, 0.5 mM dNTPs, 0.01 M DTT, 40 U of RNaseOUT, and M-MLV reverse transcriptase (Invitrogen). FastStart Universal SYBR Green Master (ROX) was used to perform quantitative RT-qPCR on a LightCycler 480 (Roche) in the presence of the following primers: ATPase, 5-CTTCAATGGGTCCCACCATA-3 (forward) and 5-CAGCAGATTTTGGCAGGTG-3 (reverse) (Esparza-Moltó et al., 2019); nuclear 18S rRNA (housekeeping gene), 5-ACGGACCAGAGCGAAAGCA-3 (forward) and 5-GACATCTAAGGGCATCACAGAC-3 (reverse) (Sheng et al., 2012). The relative levels of mRNA were calculated using the 2^−ΔΔCt^ method, where ΔCt = Ct^ATPase^ - Ct^18S^, and reported as fold change relative to PS1^control^.

### Statistical analysis

GraphPad Prism 8 (GraphPad Software Inc., USA) was employed to analyse the data. Unpaired Student’s t-test was performed to compare two independent groups. If variances were unequal, Welch’s correction was applied. If the Shapiro– Wilk test, or Kolmogorov–Smirnov test, indicated that the normality assumption was not met, the non-parametric Mann–Whitney test was used. Comparisons among more than two groups were carried out using a one-way ANOVA test followed by Tukey’s multiple comparisons post-hoc test. If the assumption of homoscedasticity was not met, Welch’s ANOVA test followed by Games–Howell multiple comparisons post-hoc test was employed. To study the effects of two independent variables on one dependent variable, a two-way ANOVA test followed by Dunnett’s post-hoc test was performed. Statistical significance was set at p ≤ 0.05 (5%), and all probabilities are reported as two-tailed.

## Acknowledgements

This study received funding from the Agencia Nacional de Promoción Científica y Tecnológica (PIBT/09-2013 to L.M. and A.C.; PICT-2015-0285 and PICT-2016-4647 to L.M.; PICT 2019-00471 and PICT 2021-00630 to C.W.).

## Data availability

All relevant data can be found within the article and its supplementary information

## Author contributions statement

Conceptualization: C.W, L.M.; Methodology: C.W., P.G., M.M.R., L.G, L.C., G.V.N. N.O; E.M; A.H.R. Statistical Analysis: C.W., P.G., M.M.R., E.M.; Resources: C.W, L.I.B, E.M.C., A.C, L.M.; Writing - original draft: C.W., P.G., M.M.R., G.V.N., L.M; Writing - review & editing: C.W., P.G., M.M.R., G.V.N., L.C, L.G, N.O., E.M., A.H.R., L.I.B., E.M.C. S.C., L.M.; Supervision: A.C., L.M.; Project administration: C.W., A.C., L.M.; Funding acquisition: C.W, A.C, L.M. All authors read and approved the final manuscript.

