## Supplementary Information for "Dysregulation of energy metabolism and calcium homeostasis in iPSC-derived neurons carrying Presenilin-1 M146L gene mutation"

**Supplementary Figures**

**Figure S1. PS1$^{M146L}$ mutation causes Familial Alzheimer`s Disease (FAD).**

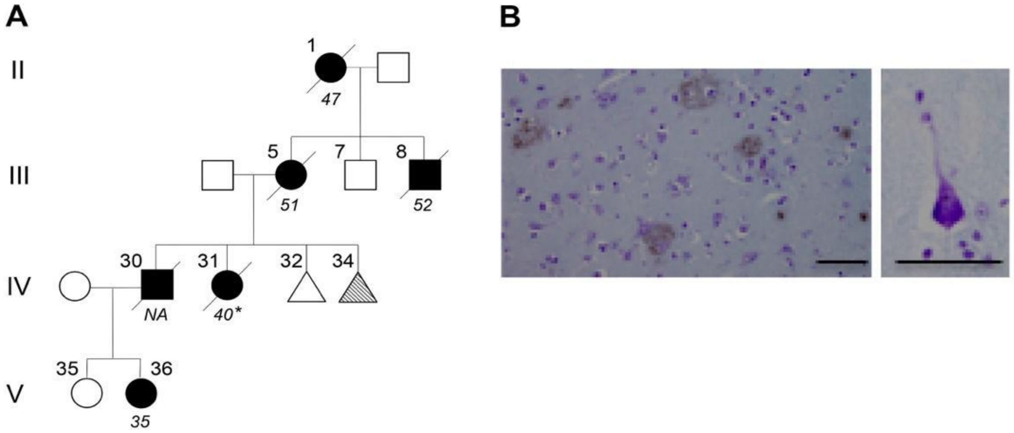


**(A)** Four-generation branch of AR1 pedigree in which early onset FAD segregates as an autosomal dominant trait. Squares, males: circles. females: filled symbols, affected; diagonal line, deceased; triangles, asymptomatic; dashed, carrier. Above each symbol, pedigree numbers; below affected individuals present ages or ages at death. Gender and age of asymptomatic members are not shown to preserve confidentiality. The asterisks indicate pathologic confirmation. Subject V-36, donant of fibroblasts to generate iPSC-derived neurons (iNs). **(B)** Pathologic confirmation in human brain of M146L carrier (IV-31). **Left panel.** Numerous senile plaques in the cortex stained with monoclonal antibody 4G8, specific for residues 17-24 of Aβ; **Right panel.** A characteristic neurofibrillary tangle stained with cresyl violet. Scale bar, 50 µm.

**Figure S2. Schematic representation of the different stages of differentiation of reprogrammed fibroblasts (iPSCs) from skin into cortical neurons.**

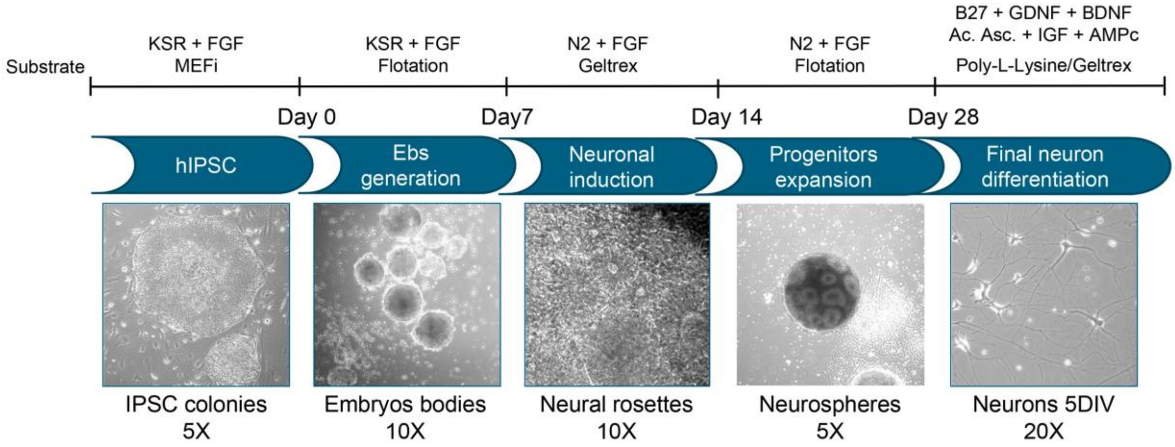


The upper panel shows the substrates used during the course of the procedure and the corresponding stage of cell characterisation. The lower panel shows phase contrast images and their corresponding magnification, showing cells in some of the most characteristic stages.

**Figure S3. Mitochondrial content is similar between PS1$^{control}$ and PS1$^{M146L}$ cells.**

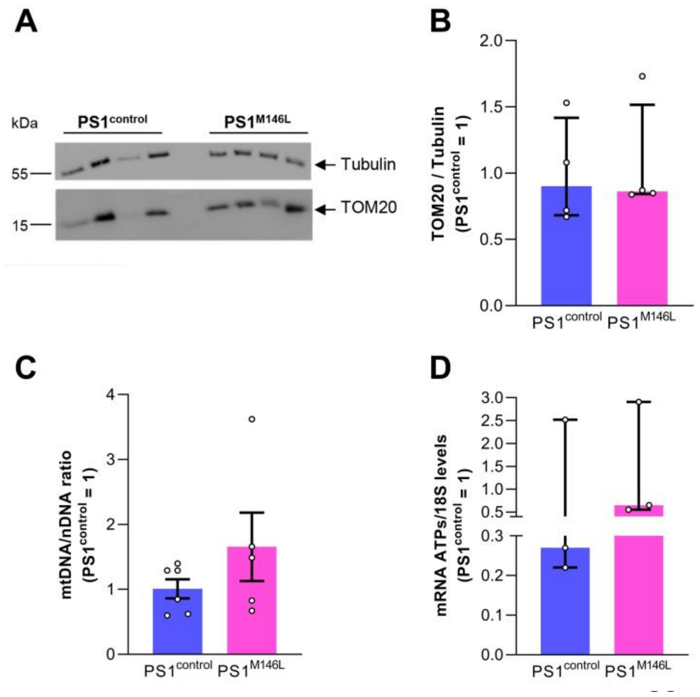


**(A)** Representative Western blot of PS1$^{control}$ and PS1$^{M146L}$ cell homogenates probed with an antibody against the mitochondrial protein TOM20 (lower panel), and Tubulin as a loading control (upper panel). **(B)** Quantification of TOM20 levels normalized to Tubulin in both genotypes (U=6, p=n.s., Mann–Whitney test). **(C)** Relative gene expression ratio of mtDNA to nDNA in both genotypes (t=1.18, df=4.63, p=n.s., Welch's correction). **(D)** mRNA transcript levels of ATP synthase (ATPs) normalized to 18S rRNA in both genotypes (U=2, p=n.s., Mann–Whitney test). Bars and error bars represent the median and interquartile range (B and D) or the mean ± SEM (C), with individual data points superimposed.

### Supplementary Tables

***Table S1. Correct mutation propagation of A>T transversion in exon 5 of clone F2A23 and F2A111***

| Clon FA111 | | | | | | | Clon F2A23 | | | | | | |
|---|---|---|---|---|---|---|---|---|---|---|---|---|---|
| **Codon** | **ref** | **total** | **A** | **C** | **T** | **G** | **Codon** | **ref** | **total** | **A** | **C** | **T** | **G** |
| **145** | G | 1937 | 0 | 1 | 2 | 1934 | **145** | G | 1663 | 1 | 1 | 0 | 1661 |
| | T | 1919 | 1 | 5 | 1913 | 0 | | T | 1650 | 1 | 3 | 1645 | 1 |
| | C | 1914 | 2 | 1910 | 2 | 0 | | C | 1647 | 3 | 1641 | 3 | 0 |
| **146** | A | 1872 | 990 | 1 | 881 | 0 | **146** | A | 1616 | 857 | 0 | 759 | 0 |
| | T | 1876 | 1 | 4 | 1871 | 0 | | T | 1644 | 0 | 2 | 1641 | 1 |
| | T | 1628 | 0 | 11 | 1617 | 0 | | T | 1644 | 0 | 1 | 2 | 1641 |

Ref, expected base; Total, total number of reads/alignments at that position. In red are shown the heterozygous A >T mutation in the first position of codon 146
